# Targeting oncogenic Wnt/β-catenin signaling in adrenocortical carcinoma disrupts ECM expression and impairs tumor growth

**DOI:** 10.1101/2022.10.23.509323

**Authors:** Morgan K. Penny, Antonio M. Lerario, Kaitlin J. Basham, Sahiti Chukkapalli, Yingjie Yu, Dipika R. Mohan, Chris LaPensee, Kimber Converso-Baran, Mark J. Hoenerhoff, Laura Suárez Fernández, Carmen González del Rey, Thomas J. Giordano, Ruolan Han, Erika A. Newman, Gary D. Hammer

**Author notes:** Co-last authors (Erika A. Newman, Gary D. Hammer).

## Abstract

Adrenocortical carcinoma (ACC) is a rare, but highly aggressive cancer with limited treatment options and poor survival for patients with advanced disease. Improved understanding of transcriptional programs engaged in ACC will help direct rational, targeted therapies. While activating mutations in Wnt/β-catenin signaling are frequently observed, the β-catenin-dependent transcriptional targets that promote tumor progression are poorly understood. To address this question, we used independent component analysis and identified a novel Wnt/β-catenin-associated signature in ACC predictive of poor survival. This signature was enriched for the extracellular matrix (ECM), suggesting a potential role for Wnt/β-catenin in regulating the ACC microenvironment. We further investigated the minor fibrillar collagen, collagen XI alpha 1 (*COL11A1*), and found that *COL11A1* expression strongly correlated with both Wnt/β-catenin activation and poor patient survival. Inhibition of constitutively active Wnt/β-catenin signaling in the human ACC cell line, NCI-H295R, significantly reduced expression of *COL11A1* and other ECM components, and decreased viability of cancer cells *in vitro*. To investigate the preclinical potential of Wnt/β-catenin inhibition *in vivo*, we developed and characterized a novel orthotopic xenograft model utilizing minimally invasive techniques. Treatment with the newly developed Wnt/β-catenin:TBL1 inhibitor Tegavivint significantly reduced tumor growth in this preclinical model. Together, our data supports that inhibition of aberrantly active Wnt/β-catenin disrupts transcriptional reprogramming of the microenvironment and reduces ACC growth and survival. Furthermore, this β-catenin-dependent oncogenic program can be therapeutically targeted with a newly developed Wnt/β-catenin inhibitor. These results show promise for further clinical development of Wnt/β-catenin inhibitors in ACC and unveil a novel Wnt/β-catenin-regulated transcriptome.

**Simple Summary:** Adrenocortical carcinoma (ACC) is a rare, often deadly cancer arising from the adrenal gland. Mortality associated with ACC remains unchanged over the last several decades. The rarity of ACC, an incomplete understanding of its molecular basis, and limited availability of pre-clinical models have hampered the development of new effective therapies. The present work aims to address these gaps with a focus on the Wnt/β-catenin cell signaling pathway, which is aberrantly activated in ~40% of ACC tumors. We discover a novel ECM program activated in ACC that is associated with Wnt/β-catenin and poor survival. Wnt/β-catenin inhibition disrupts expression of ECM genes and induces loss of cancer cell viability. To extend these findings, we develop a rapid orthotopic mouse model of ACC and demonstrate that disruption of the Wnt/β-catenin axis with novel small molecule inhibitor Tegavivint is a potential effective therapeutic strategy to reduce ACC tumor burden *in vivo*.

## Introduction

Adrenocortical carcinoma (ACC) is a rare cancer arising from the adrenal cortex. The overall 5-year survival rate for ACC is 35% (1), and is less than 10% for patients with stage 4 disease (2). Although ACC patients have more favorable outcomes with surgical resection, the majority of patients either experience tumor recurrence following surgery, or present with unresectable disease and require systemic strategies that routinely rely upon cytotoxic chemotherapies with limited benefit (2). Significant efforts have been made to develop and implement multimodal therapy but provide limited benefit, with cisplatin, etoposide, doxorubicin plus mitotane (EDP-M) providing a median progression-free survival of only 5 months (3). An improved understanding of ACC and stronger preclinical models are needed to develop rational therapies.

One molecular pathway that has been long associated with ACC is the Wnt/β-catenin pathway. This pathway is essential for development and homeostasis of many tissues including the adrenal cortex (4,5). Wnt ligand binding activates downstream signaling, leading to cytoplasmic accumulation and nuclear translocation of β-catenin. In the nucleus, β-catenin serves as a transcriptional coactivator of TCF/LEF family transcription factors to up-regulate target genes, including *AXIN2* (6), *LEF1* (7,8), and *APCDD1* (9). In the absence of pathway activation, cytoplasmic β-catenin is targeted for degradation, but in human tumors, multiple genetic alterations can result in aberrant Wnt/β-catenin activation. These include loss-of-function (LOF) mutations in negative regulators *APC* and *ZNRF3*, and gain-of-function (GOF) mutations in *CTNNB1* (the gene encoding β-catenin).

A connection between aberrant Wnt/β-catenin activation and ACC pathogenesis was first suggested by the increased incidence of adrenocortical tumors in patients with Familial Adenomatous Polyposis (FAP) (10). FAP is a hereditary cancer syndrome characterized by a germline inactivating mutation in *APC*. Enhanced Wnt/β-catenin activation was further implicated in ACC by later studies that found *CTNNB1* GOF mutations and nucleo-cytoplasmic β-catenin accumulation in a relatively high proportion of human ACC cases (11). More recently, large scale genomic studies have confirmed that genetic alterations that activate Wnt/β-catenin occur in nearly 40% of sporadic ACC cases (12,13). These include GOF mutations in *CTNNB1* (~16%), LOF alterations in *APC* (2-3%), and LOF alterations in *ZNRF3* (~20%). Moreover, increased Wnt/β-catenin activity, determined by the presence of a genetic alteration or nucleo-cytoplasmic β-catenin staining, is associated with poor prognosis in ACC patients (14,15).

Given the high prevalence of genetic alterations that enhance Wnt/β-catenin signaling in ACC, there has been considerable interest in therapeutically targeting this pathway. While upstream inhibitors that block Wnt ligand activation of cell surface receptors are attractive, activating mutations in *CTNNB1* are frequent in ACC, precluding use of such inhibitors. Strategies that directly target downstream β-catenin activity are essential to effectively block pathway activation in these tumors. While several previously developed small molecules have been shown to inhibit nuclear β-catenin binding to TCF/LEF (16), clinical development of such therapies has not been successful. Tegavivint, a recently developed inhibitor currently in clinical trial, both disrupts binding of β-catenin to transducin β-like protein 1 (TBL1 - a key adaptor protein required for β-catenin binding and transcriptional activation), and promotes SIAH-1-mediated degradation of nuclear β-catenin, which is not affected by *CTNNB1* GOF mutations at the phosphorylation site (17–20). On-target effects of Tegavivint demonstrating inhibition of Wnt/β-catenin transcriptional activity have been well-documented in various *in vitro* and *in vivo* models of acute myeloid leukemia, multiple myelomna and osteosarcoma (17,18,21,22), without overt toxicity at therapeutically effective doses (18,22).

Successfully advancing new therapeutic agents in the clinic requires *in vivo* models that faithfully recapitulate human disease. In ACC, the development of preclinical xenograft mouse models with advanced disease has proven challenging. Most approaches rely on subcutaneous, renal subcapsular, or splenic injection of tumor cells (23–26). These models require injection of millions of cells (≥ 2.5×10^6^ cells) while failing to mimic the endogenous tumor microenvironment. While genetic mouse mutants that spontaneously develop metastatic ACC have recently been described, the stochastic nature of tumor development limits the utility of such models in preclinical studies designed to test new therapeutic agents (27,28). Orthotopic injection of Wnt/β-catenin-mutated ACC cells directly into the adrenal gland provides a more biologically relevant microenvironment to support ACC growth and allows for structured analyses and comparison of pharmacologically treated to non-treated tumors. This type of approach in other cancers has proven to more closely recapitulate the biology of human tumors with respect to vascularization, chemotherapy response, and metastasis, while preserving benefits of a xenograft model including rapid and controlled tumor development capturing the heterogeneity of human cancer cells (29–31).

ECM proteins are key components of the tumor microenvironment that have been shown to play driving roles in both carcinogenesis and tumor progression (32,33). Cell-matrix interactions can drive cell anchorage, spreading, and migration; growth factor interaction by sequestration; proliferation; and differentiation (34), and abnormal ECM dynamics are a hallmark of cancer (32,35). Although Wnt/β-catenin has been shown to be an essential paracrine signaling pathway mediating adrenocortical homeostasis (5,36) and the inhibition of β-catenin activity has beern shown to inhibit growth of ACC cells in culture (37–40), a role for Wnt/ β-catenin regulating the expression of ECM components and the establishment of a tumor microenvironment has yet to be investigated.

In the present work, we identify a novel transcriptional signature of aberrant Wnt/β-catenin activation that is enriched in ECM components and associated with poor survival. We tested two independent inhibitors of Wnt/β-catenin, PKF115-584 and Tegavivint, and demonstrate that both effectively repress Wnt/β-catenin activity and coordinately inhibit *in vitro* growth of ACC cells harboring a *CTNNB1* GOF mutation. In order to translate and extend our findings *in vivo*, we implemented and characterized a novel ultrasound-guided orthotopic xenograft model and found that Tegavivint treatment significantly inhibited tumor growth in mice. Taken together, these studies support further clinical development of Wnt/β-catenin inhibitors that target β-catenin-dependent transcriptional activity in ACC.

## Materials and Methods

### Transcriptome data analysis

All analysis involving ACC-TCGA transcriptome data was performed in R (http://www.R-project.org/) using software packages from the Bioconductor portal (www.bioconductor.org). We downloaded harmonized RNA-seq counts data from the Genomic Data Commons (GDC) portal using *TCGABiolinks* (41), and performed log2-cpm normalization using *EdgeR* (42) after correcting for library size using the TMM method. We then used *MineICA* (https://rdrr.io/bioc/MineICA/) to perform Independent Component Analysis (ICA). We used the Mann-Whitney test to interrogate the association between each gene signature (component) identified by ICA and the presence of *CTNNB1* mutations by comparing the loading values of a “witness” gene (which is automatically determined by *MineICA*) in *CTNNB1-m*utated and *CTNNB1*-negative samples. We performed gene set enrichment analysis using the online *GSEA* tool (www.gsea-msigdb.org) to calculate enrichment scores using the Canonical Pathways collection from MSigDB, which is comprised of several curated gene lists from different sources including Biocarta, KEGG, and Reactome, representing a broad collection of pathways and biological processes. We used *ComplexHeatmap* (43) to generate the heatmap representing the gene signature. We used *GSVA* (Gene set variation analysis) default algorithm (44) to calculate the 5-gene canonical Wnt score as previously reported (28) and the score derived from the identified 340 genes. We used the Spearman test to calculate the correlations between selected ECM genes and the 5-gene Wnt score. We used the median 340-gene score to divide the cohort into two groups and estimate OS (overall survival) and DFS (disease-free survival) differences by the Kaplan-Meier curves and log-rank test. To divide the ACC-TCGA cohort into high- and low-expression groups according to *COL11A1* expression, we plotted the log2-CPM values of the gene as a function of the natural log-transformed hazard-ratio using smoothHR (https://github.com/arturstat/smoothHR). Using an additive Cox model, we defined the optimal cutoff as the point where the lower limit of the confidence interval of the natural log-transformed hazard ratio crossed the baseline. To generate the chromatin accessibility track from ACC-TCGA samples, we downloaded 18 bigwig files from 9 ACC samples that were generated by Corces *et al*. (45) and combined them into a single bigwig file using BigWigMerge from UCSC tools (https://github.com/ucscGenomeBrowser/kent). To generate the track with the accessibility peaks, we combined the coordinates of the corresponding peaks called by Corces et al. into a single .bed file using bedtools (https://bedtools.readthedocs.io/en/latest/index.html). To annotate these peaks, we downloaded .bed files of ChIP-seq experiments for TCF/LEF transcription factors in several different human cell lines from the UniBind database (https://unibind.uio.no). We used the JBR browser (https://github.com/JetBrains-Research/jbr) to generate the figures overlaying these tracks in the genomic regions of interest.

### *Cell culture and* in vitro *compounds*

NCI-H295R (RRID:CVCL_0458) and Y1 cell lines were obtained from American Type Culture Collection (Manassas, VA) and were cultured in a humidified incubator containing 5% CO2 at 37°C. NCI-H295R cells were grown in DMEM/Ham’s F-12 (1:1) (ThermoFisher Scientific, Waltham, MA) supplemented with 10% NuSerum I (Corning, Corning, NY), 1% Insulin-Transferrin-Selenium-Ethanolamine (ThermoFisher Scientific), and 1% penicillin/streptomycin (ThermoFisher Scientific). The human NCI-H2935R cell line has been authenticated using short tandem repeat profiling within the last three years. Y1 cells were cultured in High Glucose DMEM (ThermoFisher Scientific) supplemented with 2.5% fetal bovine serum (Sigma-Aldrich, St. Louis, MO), 7.5% horse serum (ThermoFisher Scientific), and 1% penicillin/streptomycin. All experiments were performed with mycoplasma-free cells. Tegavivint was a gift from Iterion Therapeutics, INC (Houston, TX, USA) and PKF115-584 was obtained from Tocris (Minneapolis, MN). All compounds were solubilized in DMSO.

### Assessment of cell viability

Cells were plated at a density of 300,000 cells per well in a clear 24-well plate, or 30,000 cells per well in a 96 well plate one day prior to treatment. At endpoint, cells were incubated with the colorimetric reagent alamarBlue (ThermoFisher Scientific) in accordance with manufacturer instructions. Absorbance at 570nm and 600nm was measured after 3 hours.

### qRT–PCR

RNA was extracted using RNeasy Mini Kit (QIAGEN, Hilden, Germany). Reverse transcription was performed using High Capacity cDNA Reverse Transcription Kit (Applied Biosystems) according to manufacturers instructions. The cDNA obtained was then used as template for qPCR analysis using Power SYBR Green PCR Master Mix (Applied Biosystems) and an Applied Biosystems 7300 Real-Time PCR System. Expression levels were normalized using the ΔΔCt method using *HPRT1* as a housekeeping gene. Primers used are listed in Supplementary Table 2.

### Western blots

Cells were harvested in RIPA buffer (ThermoFisher Scientific) containing cOmplete Protease Inhibitor Cocktail (Roche), PhosSTOP phosphatase inhibitors (Roche), and 50 uM ETDA, on ice for 30 min, and centrifuged at 16,000 x g for 20 min at 4°C. Protein concentrations were measured using a Pierce BCA protein assay kit (ThermoFisher Scientific), and equal amounts of total protein were separated by NuPAGE 4-12% Bis-TRIS gel in MES SDS running buffer (ThermoFisher Scientific). Protein was transferred from the gel to a nitrocellulose membrane and blocked with Odyssey Blocking Buffer (LiCor, Lincoln, NE) for 1 hour. Membranes were probed with antibodies against β-catenin (ThermoFisher Scientific MA1-2001, 1:1000), Active β-Catenin (Cell Signaling 9561 1:1500), or β-actin (Sigma-Aldrich A-5441, 1:5000) at 4°C overnight. The next day, the membrane was washed three times with TBS-T buffer, incubated with the secondary antibodies for 1 hour at room temperature, and washed three times with TBS-T and one time with PBS. Images were acquired using the LiCor Odyssey imaging system. Images were cropped to re-order samples run together on the same membrane.

### Generation of luciferase-expressing cell lines

Stable NCI-H295R and Y1 cell lines expressing pLVX-EF1α-LUC2-IRES-mCherry (46), a gift from Dr. Judith Leopold (University of Michigan, Ann Arbor, MI, USA), were generated using lentiviral transduction and purified by FACS based on mCherry expression.

### Animals and animal Care

NSG mice were housed and maintained in specific pathogen-free conditions and facilities accredited by the American Association for Accreditation of Laboratory Animal Care, and in accordance with current regulations and standards of the United States Department of Agriculture, United States Department of Health and Human Services. Animal care was overseen by the Unit for Laboratory Animal Medicine at the University of Michigan.

### Xenograft model of ACC

Intra-adrenal injections were performed on six to thirteen-week-old male immunodeficient NSG mice (The Jackson Laboratory, Bar Harbor, ME) as described previously (47). Ultrasound procedures and measurements were carried out using a Visual Sonics Vevo 2100 with a MS 550D 22-55 MHz or MS400 18-38 MHz transducer (University of Michigan Cardiovascular Center Research Core). Bioluminescent imaging was performed using an IVIS Spectrum *in vivo* imaging system (Perkin Elmer, Waltham, MA) at the Center for Molecular Imaging, University of Michigan, using previously described methods (47). For initial characterization studies, mice were euthanized after tumors exceeded 150mm^3^, 10^11^ photons/s/cm^2^/sr, 120 days post-injection, ≥15% weight loss, or all other mice in the cohort had reached endpoint. For studies testing Tegavivint treatment, 200,000 NCI-H295R cells were injected. Treatment was initiated when the tumor volume reached approximately 40-100mm^3^. Tumor volumes were extrapolated from last ultrasound using the NCI-H295R tumor doubling time of 6 days. Tegavivint was suspended in 5% dextrose, and intraperitoneal (i.p.) injection of 50mg/kg Tegavivint or vehicle was administered 5 days a week for two weeks. At necropsy, tumor volume was calculated as an ellipsoid volume = 4/3*π*(0.5*D1)*(0.5*D2)*(0.5*D3), where D1, D2, D3 are the longest measurement in the X,Y, and Z axis.

### Histopathology and immunochemistry

Tissues were fixed in 10% neutral-buffered formalin for 24 hours at room temperature, processed, paraffin embedded, and cut into 5 μm sections. Tissue characterization following routine hematoxylin and eosin staining was performed by a board-certified veterinary pathologist (MJH). For immunohistochemical staining, slides were labeled with antibodies against proCOL11A1 (DMTX1, supplied by Oncomatryx, S.L., Derio, Spain, 1:400, pH9), β-catenin (BD Biosciences 610153, 1:500, 10 mM citrate buffer pH6.3), SF-1 (Proteintech Group (PTGlabs) custom made AB_2716716, 1:1000, 10mM NaCitrate + 0.05% Tween-20 pH6), Ki67 (ThermoFisher Scientific MA5-14520, 1:200, 10mM NaCitrate + 0.05% Tween-20 pH6), or Human Nucleoli (Abcam ab190710, 1:200, 10 mM citrate buffer pH6). Images were acquired on an NIkon Optiphot-2 microscope with an Olympus DP-70 camera. Scoring methodology: proCOL11A1 and β-catenin specimens were independently assessed in a blinded fashion by two observers following these criteria: proCOL11A1 immunostaining was evaluated according to the cytoplasmatic signal, scored on a 0–3 scale, with a panel of normal and non-malignant tissues used as negative controls, and pancreatic ductal adenocarcinoma used as a positive control. For statistical analysis, tumors stained 0-2 were grouped. β-catenin immunostaining was evaluated as present or absent in the membranous, cytoplasmic, and nuclear compartments for each sample. KI67 immunolabeling was quantified using the Aperio whole slide imaging and digital analysis system. To quantify labeling, the nuclear algorithm provided was used following visual optimization and tuning to the labeling on KI67 stained slides.

For immunofluorescent staining, slides were blocked for 1 h at room temperature followed by primary antibody incubation overnight at 4°C, and primary antibodies were detected with HRP-polymer solution (Vector Laboratories) and Alexa fluor tyramide reagent (Thermo Fisher). Nuclei were counterstained with DAPI. Nonspecific staining was blocked using 2.5% horse serum for non-mouse antibodies. The M.O.M. kit (Vector Laboratories) was used for all primary mouse antibodies according to the manufacturer’s instructions. IF slides were mounted using ProLong Gold (Life Technologies) and imaged on a Nikon Optiphot-2 microscope with a CoolSNAP Myo camera.

### Statistics

Statistical analyses were performed using R, Graphpad Prism 7, or Microsoft Excel software. P-values ≤0.05 were assigned significance, and data are expressed as mean ±SD. For comparison of Kaplan-Meier survival, log-rank test was performed. For comparison between ACC cells or tumors treated with different experimental conditions, a two-tailed Student’s t-test or two-tailed Welch’s t-test (if normal distribution could not be assumed) was performed.

## Results

### A Wnt/β-catenin-driven gene signature is associated with poor patient outcomes in ACC

To identify transcriptional programs that are engaged in human ACC tumors, we analyzed RNA-seq data from 78 ACC samples included in The Cancer Genome Atlas Project (ACC-TCGA) (12). Using unsupervised ICA, we identified a signature of 340 coordinately expressed genes (Supplementary Table 1) that was significantly increased in tumors with somatic activating mutations in *CTNNB1* (p = 3.84e-7, Figure 1A). This gene signature included a spectrum of Wnt/β-catenin target genes, including *AXIN2, LEF1, NKD1, APCDD1*, and *LGR5* (Figure 1A). To investigate the biologic processes enriched in this signature, we performed gene set enrichment analysis (GSEA) using MSigDB Canonical Pathways gene set, which is comprised of several curated gene lists representing a broad collection of pathways and biological processes. Consistent with the high prevalence of *CTNNB1* mutations, nuclear β-catenin signaling was significantly enriched in this component (Figure 1B, FDR q-value = 1.03e-4).

**Figure 1.**
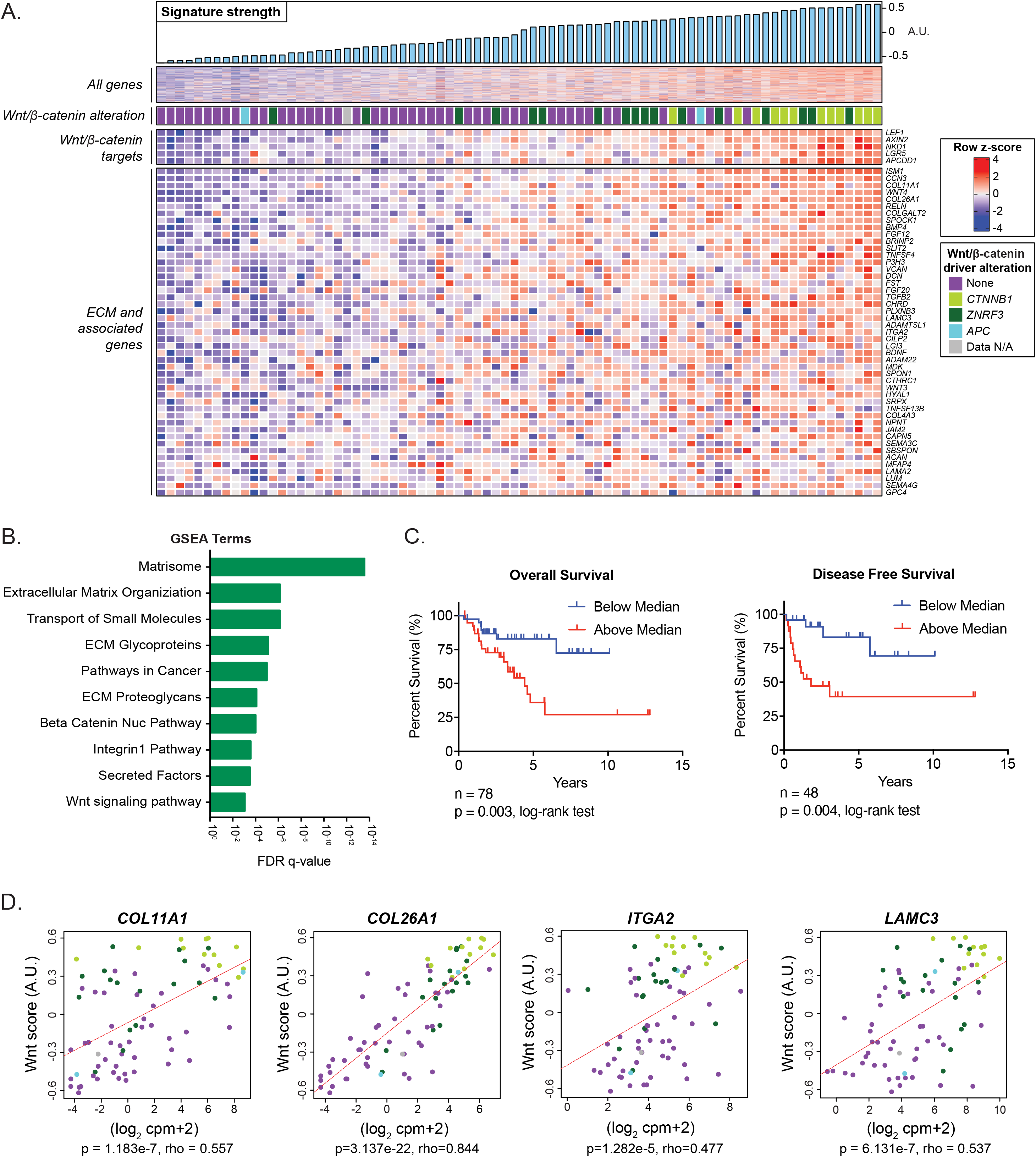
ACC-TCGA analysis identifies Wnt/β-catenin-associated ECM. A. Independent component analysis of the TCGA transcriptome dataset in ACC identifies a signature of coordinately expressed genes. A.U. stands for arbitrary units. B. GSEA identifies the signature is enriched for the Wnt/β-catenin pathway and expression of ECM and ECM-adhesion genes. C. Kaplan-Meier analysis of the TCGA cohort indicates that patients with component signature expression above the median show shorter overall survival (OS) (high score n = 39, low score n = 39). Expression higher (n = 24) or lower (n = 24) than the median also predicts shorter disease-free survival (DFS). D. Expression of *COL11A1, COL26A1, LAMC3*, and *ITGA2* is significantly correlated with a Wnt score representing expression of bona fide Wnt/β-catenin target genes *AXIN2, LEF1, APCDD1, NKD1*, and *LGR5*.

Given the previously reported associations between Wnt/β-catenin pathway activation and poor outcomes in ACC (14,15), we sought to determine whether this association was also captured by the identified 340 gene signature. We used *GSVA* to calculate a score that quantifies expression of these 340 genes in each sample. We divided the ACC-TCGA cohort into two groups according to the median of the 340-gene score. We observed a significant difference in both overall survival and disease-free survival between the two groups (OS p=0.003; DFS p=0.004) (Figure 1C). These results support that aberrant Wnt/β-catenin activation is associated with poor patient outcomes in ACC, consistent with prior studies (14,15).

### Wnt/β-catenin activity is associated with ECM expression

In addition to these findings, GSEA revealed that the Wnt/β-catenin-associated signature was most heavily enriched for ECM, ECM-receptor proteins, and other ECM-associated proteins, including collagens, integrins, laminins, and secreted factors (Figure 1B). Given the strong association between this gene signature and *CTNNB1* mutation, we hypothesized that the signature, in which ECM biology is among the most significantly represented processes, was Wnt/β-catenin-driven. To further investigate this hypothesis, we performed correlation analysis of select ECM genes and known Wnt/β-catenin target genes. We analyzed ECM genes that provided the highest component projections in the signature identified by ICA, in the class of collagen, laminin, and integrin: *COL11A1, COL26A1, LAMC3*, and *ITGA2* (Supplementary Table 1). To more accurately measure Wnt/β-catenin pathway activation and not rely on a single gene readout, we calculated a Wnt score based on the combined expression of 5 bona fide Wnt/β-catenin target genes (28). Our analysis showed a significant positive correlation between the Wnt score and each of the ECM genes (Figure 1D). Notably, tumors with *CTNNB1* mutations exhibited the highest Wnt scores and the highest expression of ECM genes. Additional analysis using publicly available datasets demonstrated TCF/LEF binding in the promoter and/or putative distal regulatory elements of our selected subset of ECM genes, overlapping with regions of accessible chromatin in ACC ATAC-seq data (Supplementary Figure 1).

High expression of minor fibrillar collagen *COL11A1* has been associated with disease progression and poor survival in ovarian and other cancers (48). We therefore chose to investigate expression of this gene further. We analyzed patient data from ACC-TCGA and found that high *COL11A1* transcript expression was associated with significantly shorter OS and DFS (Figure 2A, Supplementary Figure 2). To follow up on these findings, we performed immunohistochemistry on tissue microarrays (TMAs) containing 97 ACC samples. Recent work has determined that ACC has one of the lowest contributions of stromal cells when compared to other cancer types (12), suggesting that ECM may be at least in part tumor-cell-derived. In the current study, immunohistochemistry showed COL11A1 expression in the tumor parenchyma of ACC. Moreover, we observed that COL11A1 staining was significantly enriched in tumors with nuclear β-catenin localization (p = 0.006) (Figure 2B), indicating that expression of COL11A1 protein is associated with oncogenic Wnt/β-catenin activation. Outcomes data associated with TMA of ACC samples confirmed that COL11A1 protein was also significantly associated with shorter OS (p=0.0003) and DFS (p=0.0075) (Figure 2C). Staining of tumors from the NCI-H295R human ACC cell line, which harbors an activating S45P *CTNNB1* mutation (11), found co-expression of the steroidogenic marker SF-1 and COL11A1, indicating that COL11A1 is expressed by ACC cells (Figure 2D).

**Figure 2.**
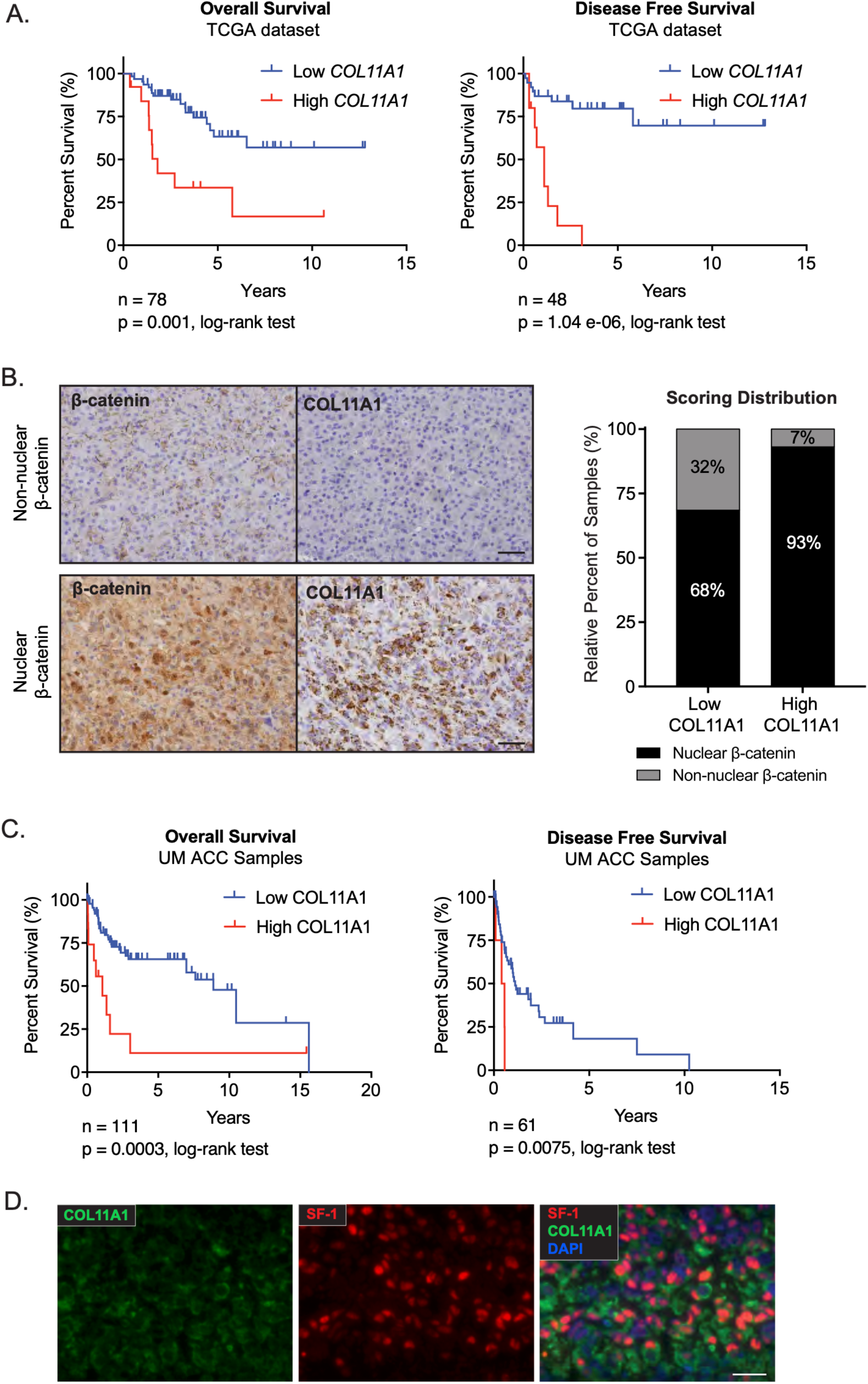
COL11A1 expression is correlated with Wnt/β-catenin activation in human ACC samples and predicts outcome. A. Correlation of OS and DFS with *COL11A1* expression by Kaplan–Meier analysis in TCGA datasets shows high *COL11A1* transcript expression is correlated with decreased OS (n=66 low, n=12 high, log-rank test p=0.001) and DFS (n=38 low, n=10 high, log-rank test p=1.04e-06). B. Serial sections from patient ACC samples (n=97) collected at the University of Michigan and stained for β-catenin and COL11A1 show that COL11A1 expression is correlated with nuclear β-catenin localization (binomial proportions test p = 0.006). Representative serial sections shown from a tumor with non-nuclear β-catenin localization versus nuclear β-catenin localization. C. Kaplan-Meier analysis of OS and DFS following COL11A1 staining of University of Michigan ACC samples shows high COL11A1 staining is correlated with shorter OS (n=98 low n=13 high, log-rank test p=0.0003) and shorter DFS (n=56 low, n=5 high, log-rank test p=0.0075). D. NCI-H295R xenograft tumor with COL11A1 and SF-1 immunofluorescent staining demonstrates co-expression in ACC cells. Scalebar, 100 μM.

### Inhibition of Wnt/β-catenin reduces ACC viability and disrupts ECM expression

Having established that Wnt/β-catenin activity and *COL11A1* expression were correlated with poor outcomes in patients, we next wanted to evaluate the effects of Wnt/β-catenin inhibition on the ECM gene signature and investigate the utility of inhibition as a therapeutic strategy in ACC. We performed experiments on the NCI-H295R human ACC cell line, which demonstrates high levels of Wnt/β-catenin activity compared to Y1 ACC cells (wild type β-catenin) (Supplementary Figure 3A). NCI-H295R cells were treated with Tegavivint, a recently developed Wnt/β-catenin inhibitor in clinical development (17,21) or PKF115-584, a well-established Wnt/β-catenin inhibitor (16) previously shown to inhibit cell growth *in vitro* in ACC (37). Both inhibitors caused a dose-dependent inhibition of cell growth and viability (Figure 3A, 3B) with NCI-H295R cells showing greater sensitivity than Y1 cells (Supplementary Figure 3B). Given that Tegavivint causes a decrease in β-catenin levels (17,18), we investigated the effect of treatment in NCI-H295R cells and found that Tegavivint resulted in a robust decrease of GOF β-catenin protein (Figure 3C).

**Figure 3.**
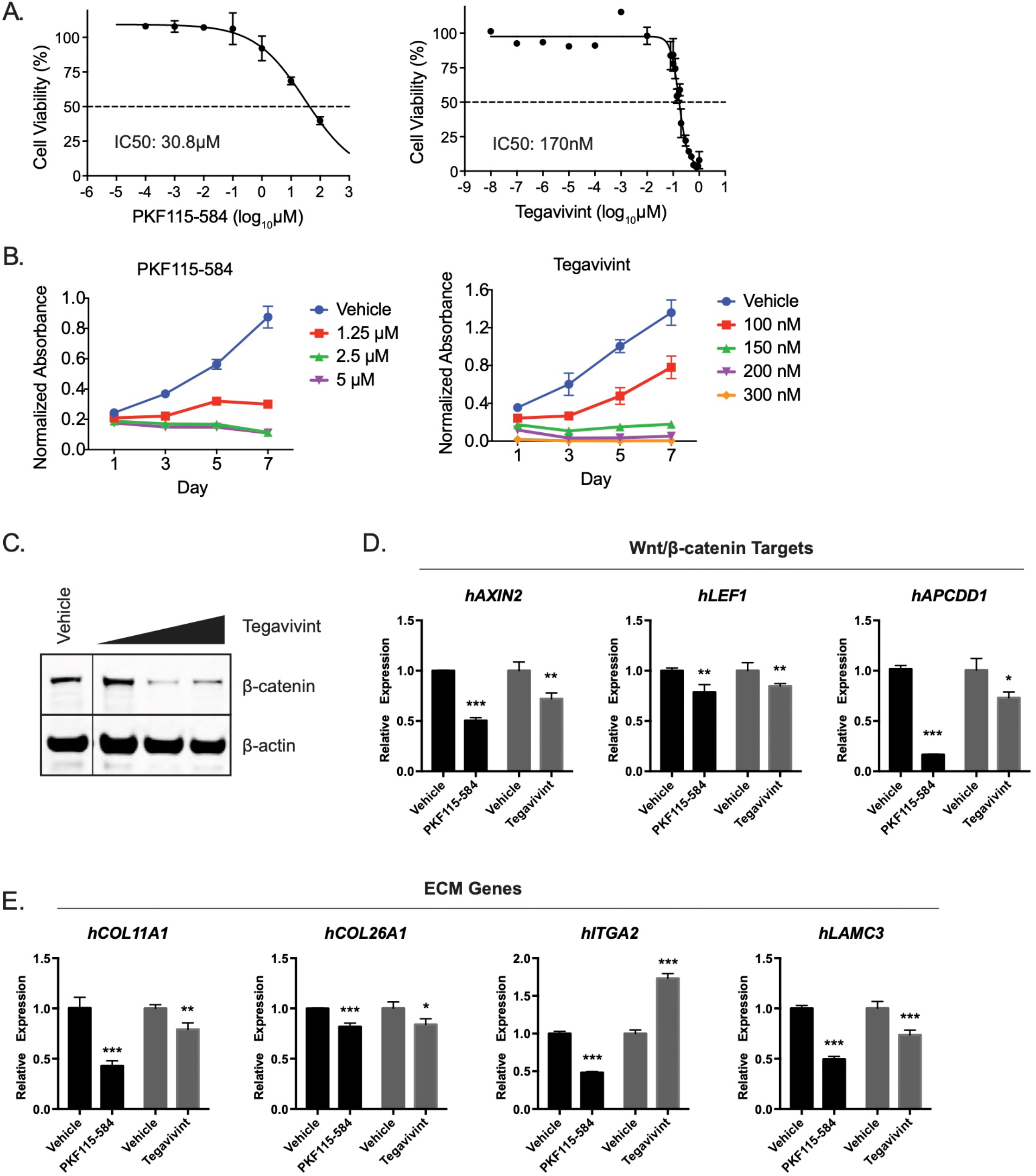
Constitutively Wnt/β-catenin-active ACC cells are sensitive to β-catenin inhibition *in vitro*. A. IC50 for NCI-H295R ACC cells treated with PKF115-584 or Tegavivint for 24 hours B. β-catenin inhibition with PKF115-584 or Tegavivint treatment leads to significantly decreased NCI-H295R viability, suggesting Wnt/β-catenin signaling may regulate cell viability in ACC. C. Representative western blot following 24hr treatment with 100 nM, 150 nM, or 200 nM Tegavivint. Images were cropped to re-order samples run together on the same membrane. D. Gene expression of Wnt/β-catenin targets in NCI-H295R cells following 24 hour treatment with 100 nM Tegavivint or 1.25 μM PKF115-584. E. Gene expression of Wnt/β-catenin-associated ECM in NCI-H295R cells, following 24 hour treatment with 100 nM Tegavivint or 1.25 μM PKF115-584. p-value was calculated using two-tailed Student’s t-test. Data are presented as mean□±□SD; *p<0.05; **p<0.005, ***p<0.0005.

To specifically interrogate the link between Wnt/β-catenin transcriptional activity and ECM expression in ACC, we treated NCI-H295R cells with Tegavivint and PKF115-584, and measured gene expression changes by qRT-PCR. Both inhibitors effectively blocked Wnt/β-catenin activation, as evidenced by significantly reduced expression of *AXIN2, LEF1, and APCDD1* (Figure 3D). Moreover, Wnt/β-catenin inhibition significantly decreased *COL11A1, COL26A1*, and *LAMC3* expression (Figure 3E). These results demonstrate that pharmacological disruption of Wnt/β-catenin activity in ACC also decreases expression of ECM components.

### An orthotopic xenograft model of ACC demonstrates high-grade ACC with metastatic potential

We next wanted to test the efficacy of Wnt/β-catenin inhibition *in vivo*. Preclinical models of ACC are limited and existing heterotopic xenografts, including subcutaneous models, do not mimic the endogenous tumor microenvironment (23,25,26). Development and use of orthotopic xenografts, however, has been limited by complex and morbid murine surgery (27,34). To overcome these limitations, we established and characterized a novel orthotopic ACC xenograft model utilizing ultrasound-guided implantation of tumor cells (47).

First, we generated two stable ACC cell lines, NCI-H295R and Y1 cells, expressing a luciferase construct for *in vivo* visualization (46). We then used ultrasound guidance to inject 200,000 Y1 or NCI-H295R cells in the left adrenal gland of NSG immunocompromised mice (Figure 4A). An additional group of animals was injected with 1,000,000 NCI-H295R cells since NCI-H295R xenografts have previously required millions of cells to achieve efficient tumor engraftment (23). Following implantation, we monitored mice for tumor growth. Because tumor growth in the retroperitoneal space cannot be measured directly, we used ultrasound with 3D reconstruction of tumor boundaries as well as bioluminescent imaging to characterize tumor engraftment and progression (Figure 4B). All groups demonstrated ≥80% engraftment efficiency (Table 1).

**Figure 4.**
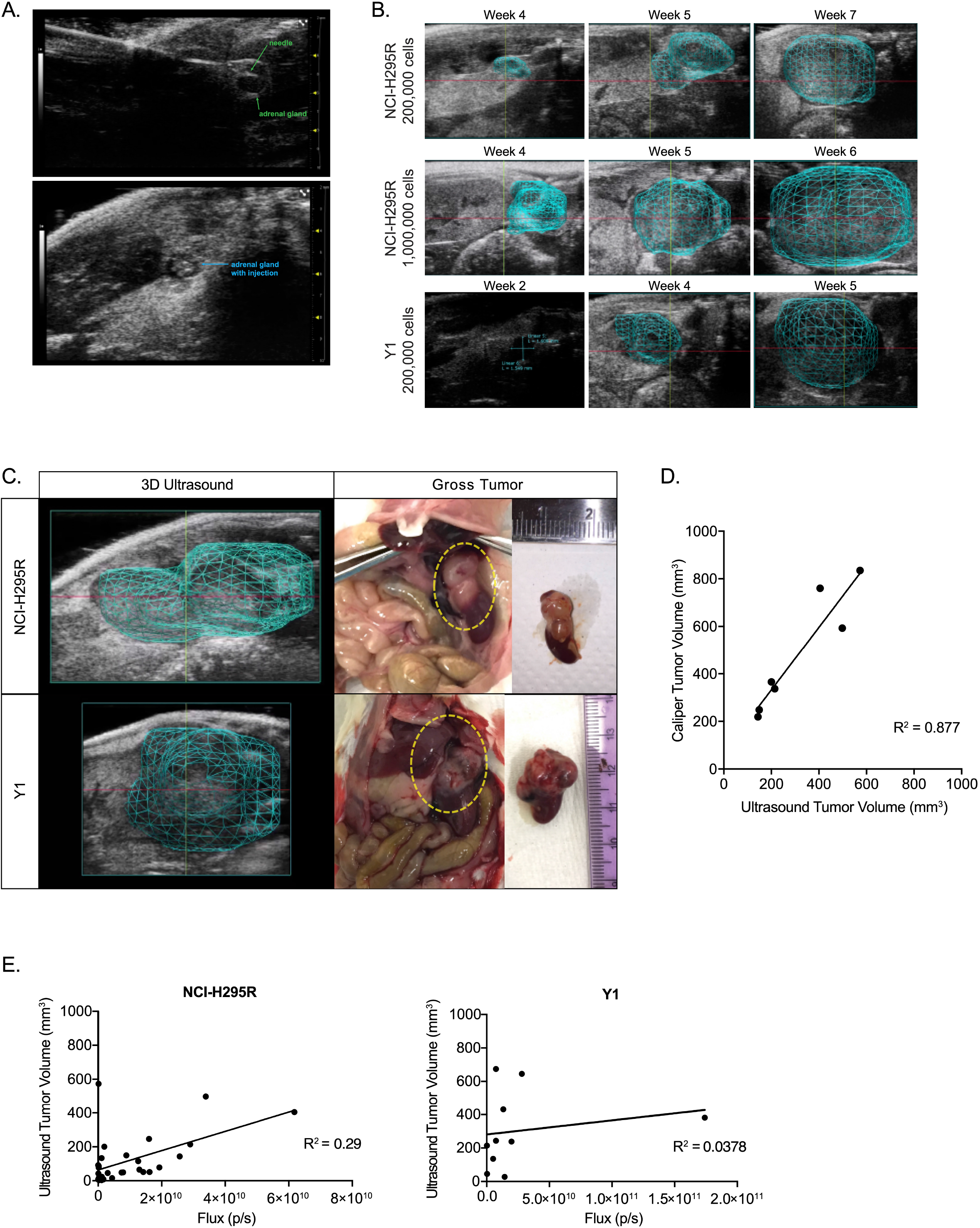
Xenograft implantation and tumor progression. A. *In vivo* injection process B. Adrenals and the growth of tumors were monitored using ultrasound, with 3D reconstruction of tumor boundaries when tumor is detected C. Here, tumor boundaries mapped from ultrasound prior to necropsy are shown alongside gross tumor. D. Ultrasound tumor volume and tumor volume calculated from caliper measurements are strongly correlated (linear regression R^2^ = 0.877). E. *In vivo* bioluminescent signal is plotted against ultrasound tumor volume (NCI-H295R linear regression R^2^ = 0.29; Y1 linear regression R^2^ = 0.0378).

**Table 1.** Tumor characteristics following orthotopic xenograft implantation.

| Cell Line | Cell Number | Average Time to Detection (SD) | Average Time to Endpoint (SD) | Tumor Engraftment | Liver Metastases | Lung Metastases |
| --- | --- | --- | --- | --- | --- | --- |
| Y1 | $2 \times 10^5$ | 2.5 weeks (0.9) | 5.1 weeks (1.0) | 89% | 50% | 83% |
| NCI-H295R | $2 \times 10^5$ | 3.6 weeks (0.9) | 8.9 weeks (3.0) | 80% | 0% | 0% |
| | $1 \times 10^6$ | 3.4 weeks (0.7) | 6.9 weeks (2.7) | 100% | 13% | 0% |

To validate the accuracy of ultrasound tumor volume reconstruction, we compared ultrasound measurements taken just prior to necropsy with the gross tumor specimen. We found that 3D ultrasound tumor volume was a sensitive and accurate measurement of tumor morphology (Figure 4C) that strongly correlated with tumor volume calculated from caliper measurements at necropsy (R^2^ = 0.877) (Figure 4D). We also compared bioluminescent signal to ultrasound volume and found that total bioluminescent flux (photons/second) measured *in vivo* did not correlate linearly with tumor volume (Figure 4E). Taken together, these data supports 3D ultrasound as a more accurate and preferred method for adrenal tumor monitoring.

At necropsy, tumors were characterized to identify utility and timing of the model for future studies. Histologically, tumors formed from NCI-H295R and Y1 xenografts modeled characteristics of high-grade ACC, designated by 20 or greater mitotic figures per 50 high power fields (≥20/50 hpf). NCI-H295R tumors formed following injection with 200,000 cells and 1,000,000 cells had >20/50 hpf (mean 582.5± 79.3 SD; mean 570±133.3 SD), and a high Ki67 labeling index (Figure 5B, 5G). All Y1 tumors had >20/50 hpf (mean 346.7 ± 228.1 SD), and a high Ki67 labeling index (Figure 5E, 5G). Tumors also retained expression of adrenocortical marker steroidogenic factor-1 (SF-1) (Figure 5C, 5F). Importantly, xenografted tumors formed distant metastases in liver and lung, sites characteristic of ACC (Table 1). While NCI-H295R metastases were found in the liver in one mouse with long tumor latency (Figure 5H, 5I; Table 1), Y1 metastases frequently formed in the liver and lung (Figure 5J, 5K; Table 1). Our characterization establishes that orthotopic adrenal xenografts model high-grade, advanced ACC.

**Figure 5.**
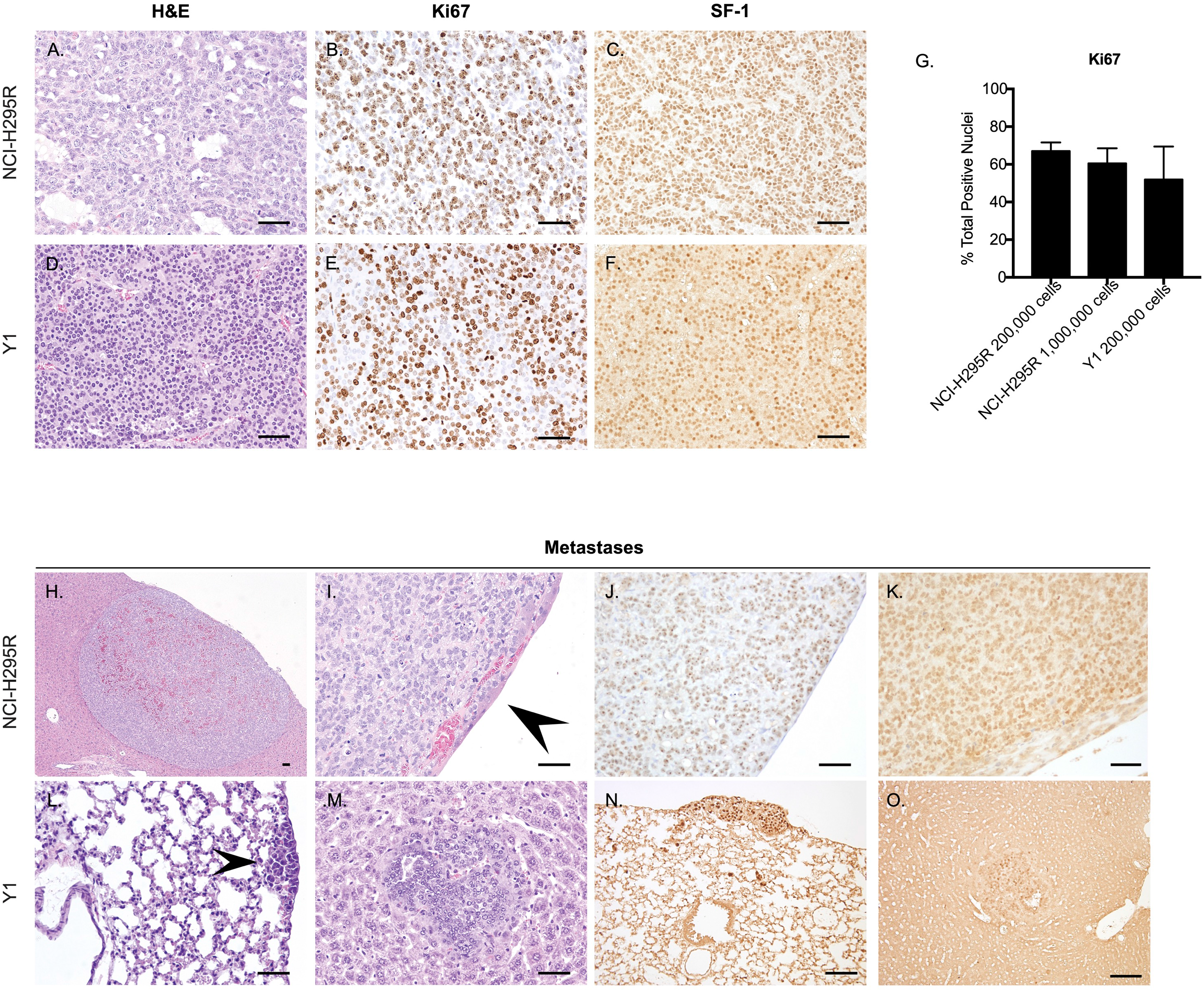
Histology of primary tumors and metastatic growths. A. NCI-H295R tumors were characterized as composed of an expansile and focally infiltrative proliferation of packets, clusters, and sheets of neoplastic epithelial cells interspersed with variably sized cystic spaces. B. NCI-H295R Ki67 staining and C. SF-1 staining. D. Tumors formed from Y1 cells were composed of ribbons, clusters, and lobules of poorly differentiated epithelial cells separated by a fine fibrovascular stroma, with multifocal areas of hemorrhage and necrosis. E. Y1 Ki67 staining and F. SF-1 staining. G. Ki67 quantification (mean□±□SD). H.,I. NCI-H295R liver metastasis, overlain with a layer of hepatocytes. J. Anti-human Nucleolar staining K. SF-1 L. Y1 lung metastasis. M. Y1 liver metastasis. N., O. SF-1 staining of Y1 metastases. Scalebar, 100 μM.

### *Tegavivint inhibits high-grade ACC growth* in vivo

To investigate therapeutic treatment of ACC, we next evaluated the effect of β-catenin inhibition on tumor growth in our xenograft model. To best model therapeutic treatment in patients with existing ACCs, NCI-H295R cells, which harbor an activating GOF β-catenin mutation, were implanted. Tumors were allowed to grow to 40-100mm^3^ after which they were treated with 50 mg/kg Tegavivint, or vehicle (Figure 6A). There was no significant difference in estimated tumor volume at the onset of treatment (p = 0.936) and Tegavivint was well tolerated by mice. After two weeks of Tegavivint treatment, a striking 65% reduction in tumor weight and 58% reduction in tumor volume was observed over vehicle-treated controls (p=0.0047, p=0.0015, respectively; Figure 6B). Though tumors contained extensive areas of necrosis, staining of viable areas showed that Tegavivint produced a mild reduction in β-catenin levels (Supplementary Figure 4). Taken together, we show that Wnt/β-catenin inhibition is a potentially efficacious therapeutic strategy for high-grade ACC harboring activating mutations in Wnt/β-catenin signaling.

**Figure 6.**
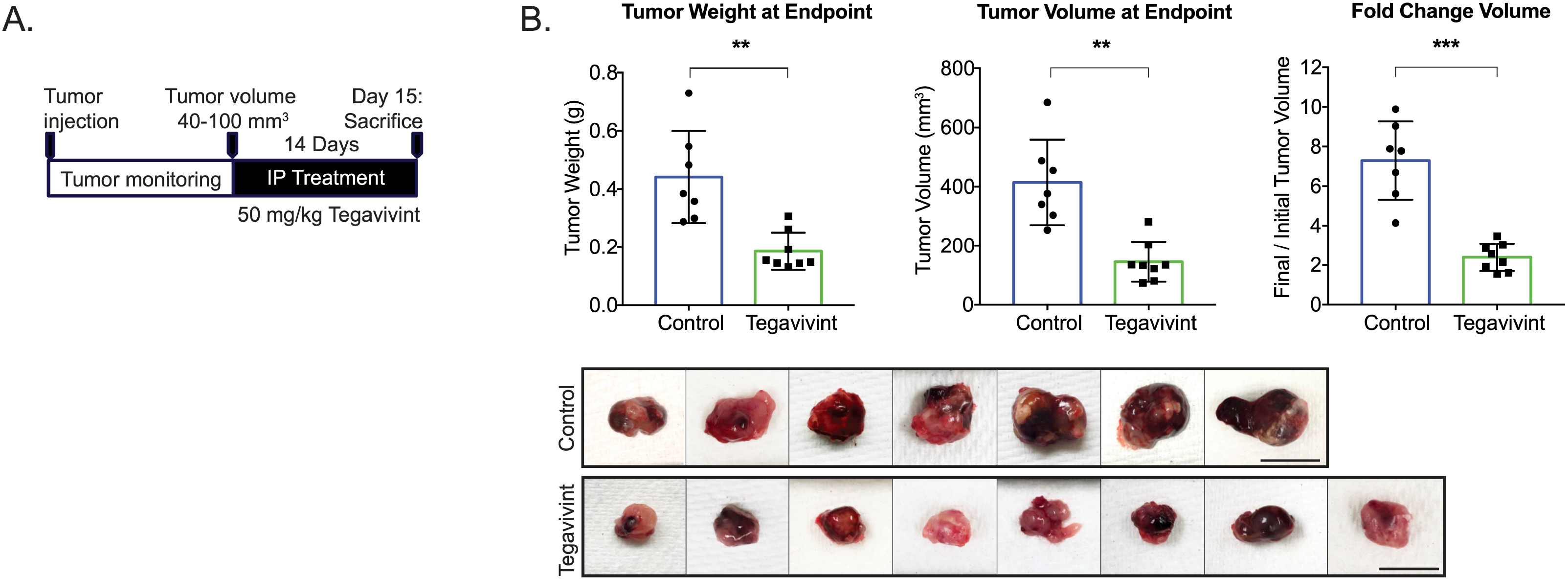
β-catenin inhibition significantly reduces tumor growth. A. Experimental scheme of study design. NSG mice were injected with 200,000 NCI-H295R cells in the left adrenal. Treatment was started when xenografts reached 40-100mm^3^. Mice were treated with 50 mg/kg Tegavivint or vehicle 5 days per week (Monday - Friday) for 14 days. B. Tegavivint treatment significantly decreased tumor weight and volume at endpoint. p-value was calculated using two-tailed Welch’s t-test. Data are presented as mean□±□SD; *p<0.05; **p<0.005, ***p<0.0005. Scalebar, 10 mm.

## Discussion

Over the last two decades, a large body of evidence has implicated sustained Wnt/β-catenin signaling as a key driver event in the molecular pathogenesis of ACC (50). Indeed, we now know that somatic alterations targeting Wnt/β-catenin pathway prevail in nearly 40% of ACC (12,13). Furthermore, genetic mouse models bearing adrenal-specific activation of Wnt/β-catenin exhibit adrenocortical hyperplasia, dysplasia, and tumor formation (14,28,51). While these studies strongly support a dominant role for Wnt/β-catenin in the molecular pathogenesis of ACC, much is unknown about the downstream mechanisms by which Wnt/β-catenin promotes tumor growth. Using ICA, we characterized a strong effect of Wnt/β-catenin activity on the ACC transcriptome. Our data builds on previous studies (14,38) to identify a 340-gene signature expressed in ACCs with *CTNNB1* mutations that is highly correlated with both Wnt/β-catenin activity and is uniquely associated with ECM expression. The strong transcriptional signature associated with the activation of this pathway and its major contribution to decreased patient survival suggest that pharmacologically targeting Wnt/β-catenin is an attractive therapeutic approach that can fulfill the urgent need to improve standard care for advanced ACC (2,3). Indeed, we show here that the transcriptional and tumor viability programs coordinated by Wnt/β-catenin in ACC are highly sensitive to pharmacological inhibition of this pathway.

Our analysis sought to better understand the role of Wnt/β-catenin signaling in ACC. Interestingly, we see a spectrum of Wnt/β-catenin signaling across ACC, with patient tumors harboring activating β-catenin mutations clustering at the highest level of signaling, whereas tumors with ligand-dependent signaling alterations in ZNRF3 show less intense Wnt/β-catenin activation, as previously observed (28). ACCs with ligand-dependent signaling demonstrate a spectrum of Wnt target gene expression. Furthermore, we observe expression of *COL11A1, COL26A1, LAMC3*, and *ITGA2*, is linearly correlated with pathway activation (Figure 1D). Though our analysis of TCGA data is limited by the absence of normal adrenal tissue, we hypothesize that only highly active Wnt/β-catenin signaling can induce expression of select ECM genes in ACC, driving a cancer-specific tumor microenvironment.

Though mutation data is not available for the TMA tumors we characterized, we expect nuclear β-catenin staining captures tumors with high Wnt/β-catenin activity, including those with *CTNNB1* and *ZNRF3* mutations, ultimately compressing the dynamic range of Wnt/β-catenin signaling by classifying tumors with both mutations (and therefore different levels of Wnt/β-catenin activation) in the same strata. IHC shows nearly all ACCs exhibiting COL11A1 protein expression have nuclear β-catenin. Such an observation is consistent with nuclear, or active β-catenin being necessary but not sufficient, to induce COL11A1 expression as measured by IHC. While it is unclear what other factors influence this relationship, the strength of Wnt/β-catenin signaling is a critical factor in determining the expression of COL11A1 in ACC.

We speculate that the correlation between Wnt/β-catenin and ECM is more broadly applicable. Indeed, differential ECM secretion has been linked to Wnt/β-catenin signaling in Ewing Sarcoma cells (52). While our results support that the Wnt/β-catenin-regulated transcriptome in ACC is heavily enriched for ECM, and demonstrate that COL11A1 accumulates in ACC samples with nuclear β-catenin staining, it is predicted that additional signaling pathways can influence ECM gene expression as well. We hypothesize that this is likely the case for ITGA2, where we observed variable response to Wnt/β-catenin inhibitors.

In other studies, COL11A1 expression has been almost exclusively localized to cancer-associated stromal cells (48,53). In contrast, ACC is one of the most stroma-poor cancer types in the TCGA cohort (12) and we found that *COL11A1* expression is present in both NCI-H295R cells and cancer cells in human samples. Though stromal contribution to ECM cannot be ruled out, our studies support that ACC cells provide a biologic contribution to the tumor microenvironment.

Our finding that *COL11A1* transcript and protein expression is associated with shortened OS and DFS is consistent with findings in ovarian cancer (44), and is complementary to findings that elevated *COL11A1* expression is associated with chemoresistance in human tumors (36,50). These results pose the intriguing question of whether ECM and COL11A1 specifically plays a causative role promoting ACC aggressiveness and progression.

Given our desire to ultimately target therapy against the Wnt/β-catenin pathway, we developed a model to translate ACC *in vitro* findings *in vivo*. We implemented and characterized a novel model of ACC that captures features of high-grade ACC in patients. The high mitotic rate and Ki67 labelling index of NCI-H295R and Y1 tumors capture histologic features of patient tumors with high risk for recurrence and decreased overall survival (2,56). We also show that orthotopic adrenal implantation can lead to spontaneous metastatic growth in the liver and the lungs, which are the most frequent sites of ACC metastasis in patients (1). Therefore, this study represents the first reported model of high-grade, spontaneously metastatic ACC using currently available cell lines. Though we did not compare orthotopic xenografts to subcutaneous or other xenografts in a controlled study, orthotopic implantation allows for modeling the microenvironment within the anatomic location of primary tumor initiation. Traditionally, intra-adrenal implantation has lacked implementation for preclinical therapeutic investigations because of the extensive surgical procedure and post-surgical monitoring (49). In contrast, our approach employing minimally invasive, ultrasound-guided implantation enabled high-throughput injection procedures, efficient tumor engraftment, and rapid recovery. We believe that the methodologies presented here will enable broader implementation of orthotopic xenografts in ACC research.

We used our preclinical tumor model to extend our *in vitro* findings and confirm therapeutic inhibition of Wnt/β-catenin significantly inhibits growth of tumors harboring a GOF *CTNNB1* mutation. Our results are consistent with and complimentary to prior studies in which β-catenin knockdown reduced Wnt/β-catenin transcriptional activity in ACC cells and impaired the function of β-catenin in cell-cell adhesion (36,39). Taken together, these data demonstrate preclinical support for further study on the clinical utility of Wnt/β-catenin inhibitors in ACC. Currently, Tegavivint is undergoing clinical testing (NCT04780568, NCT04874480, NCT04851119) in a variety of Wnt/β-catenin active cancers. It will be important to test the effectiveness of Wnt/β-catenin inhibitors in ACC as single agents or in combination with other therapeutic modalities. Our model complements findings from a recently developed genetic mouse model of β-catenin GOF with p53 loss of function which represents an alternate approach for testing further therapies directed against Wnt/β-catenin (27).

In conclusion, we have identified that ACC with high Wnt/β-catenin activity exhibits a unique expression pattern of ECM components that furthermore is associated with poor survival. Moreover, Wnt/β-catenin inhibition disrupts ECM expression and reduces growth in ACC with an activating *CTNNB1* mutation. Importantly, we developed a novel orthotopic xenograft model to investigate the preclinical implications of these findings and found that *in vivo* inhibition of Wnt/β-catenin:TBL1 signaling with Tegavivint significantly reduced tumor growth. Collectively, our studies demonstrate preclinical efficacy of a Wnt/β-catenin inhibitor in ACC, and underscore the rationale for therapeutically translating Wnt/β-catenin inhibition in ACC patients.

## Abbreviations

ACC: Adrenocortical carcinoma
DFS: Disease-free survival
ECM: Extracellular matrix
FAP: Familial Adenomatous Polyposis
GSEA: Gene set enrichment analysis
GOF: Gain-of-function
GSVA: Gene set variation analysis
HPF: high power fields
ICA: Independent Component Analysis
OS: Overall Survival
LOF: Loss-of-function
SD: Standard Deviation
TCGA: The Cancer Genome Atlas Project
TMA: Tissue Microarray

## Acknowledgements

Tegavivint was a gift from Iterion Therapeutics, INC., made possible in part by funding of Iterion Therapeutics (fka Beta Cat Pharmaceuticals) through the Product Development Award CP130058 from the Cancer Prevention and Research Institute of Texas (CPRIT). We are grateful to Dr. Judith Leopold for the pLVX-EF1α-LUC2-IRES-mCherry vector.

## Author Contributions

Conceptualization, Morgan K. Penny and Gary D. Hammer; Methodology, Morgan K. Penny, Kaitlin J. Basham, Sahiti Chukkapalli, Yingjie Yu, Dipika R. Mohan, Chris LaPensee, Kimber Converso-Baran, Mark J. Hoenerhoff, Laura Suárez Fernández, Carmen González del Rey, Ruolan Han, Erika A. Newman; Data Analysis, Morgan K. Penny, Antonio M. Lerario; Validation, Morgan K. Penny, Chris LaPensee, Dipika R. Mohan; Investigation, Thomas J. Giordano, Erika A. Newman, Gary D. Hammer; Resources, Laura Suárez Fernández, Carmen González del Rey, Ruolan Han, Thomas J. Giordano, Erika A. Newman, Gary D. Hammer; Writing – Original Draft Preparation, Morgan K. Penny; Writing – Review & Editing, Antonio M. Lerario, Kaitlin J. Basham, Chris LaPensee; Visualization, Morgan K. Penny and Antonio M. Lerario; Funding Acquisition, Erika A. Newman, Gary D. Hammer

## Institutional Review Board Statement

Studies involving human tissue samples were approved by the University of Michigan Institutional Review Board (HUM00045656). The samples consisted of formalin-fixed, paraffin-embedded tissues from 97 patients embedded in TMAs. All animal care and animal experimental procedures were performed in accordance with the University Committee on Use and Care of Animals at the University of Michigan (PRO00008248).

## Data Availability Statement

Data sources and handling of the publicly available datasets used in this study are described in the Materials and Methods. Further details and other data that supports the findings of this study are available from the corresponding authors upon request

## Supplementary Figure Legends

**Supplementary Figure 1.** TCF/LEF binding in promoters and/or putative distal regulatory elements of our selected subset of ECM genes, from publicly available datasets overlapping with accessible chromatin in ACC ATAC-seq data. ChIP-seq datasets are human cell lines. ACC ATAC-seq open chromatin overlap with TCF/LEF peaks are highlighted in pink.

**Supplementary Figure 2.** Regression analysis of *COL11A1* expression and A. OS hazard ratio or B. DFS hazard ratio from ACC-TCGA data.

**Supplementary Figure 3. ACC cell line Wnt/β-catenin signaling and sensitivity to β-catenin inhibition *in vitro*.**

A. Western blot of untreated cells from NCI-H295R, Y1, and ATC7L ACC cell lines. B. IC50 for Y1 cells treated with Tegavivint or PKF115-584 for 24 hours. C. NCI-H295R viability following treatment with Tegavivint or PKF115-584 for 24 hours. Data are presented as mean□±□SD.

**Supplementary Figure 4. Effects of β-catenin inhibition *in vivo*.**

A. Murine ACC tumors treated with 50 mg/kg Tegavivint for 2 days and stained for β-catenin. Scalebar, 100 μM.

