## Supplementary figures and images for "Targeting oncogenic Wnt/β-catenin signaling in adrenocortical carcinoma disrupts ECM expression and impairs tumor growth"

### Supplementary Figure 1

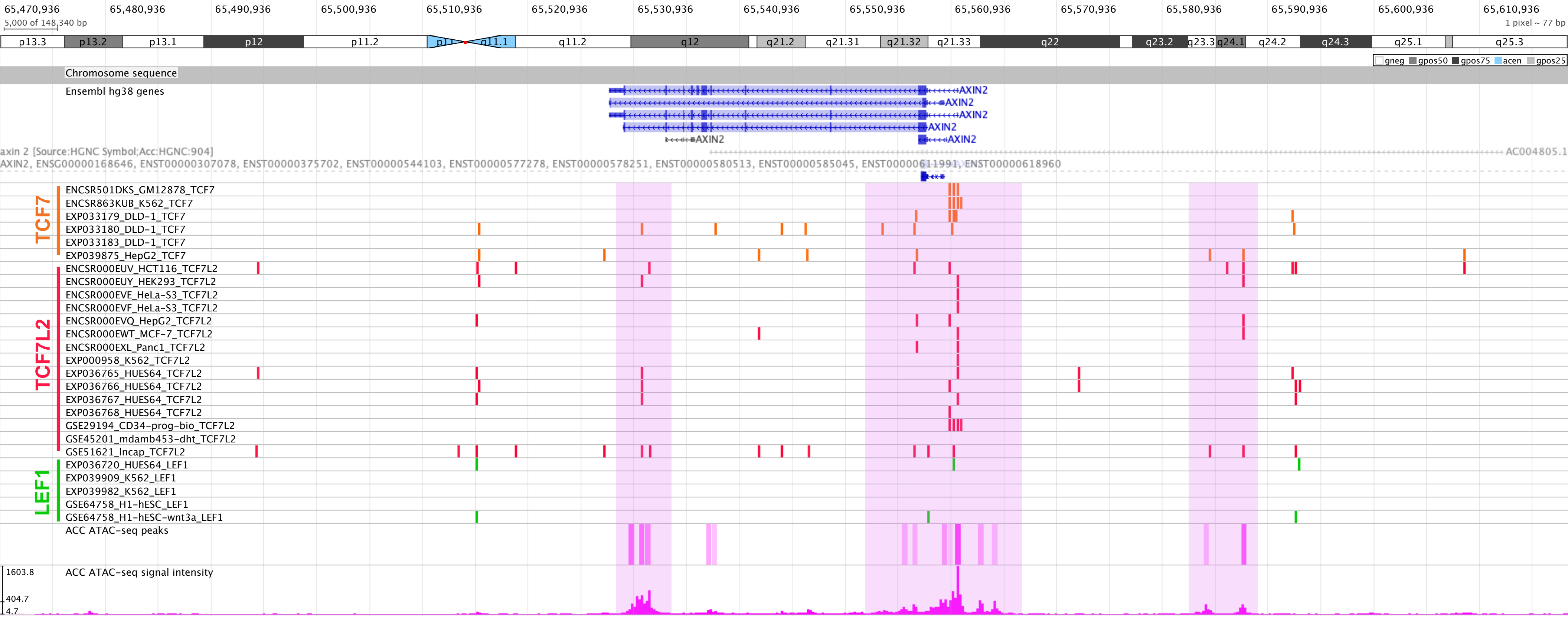

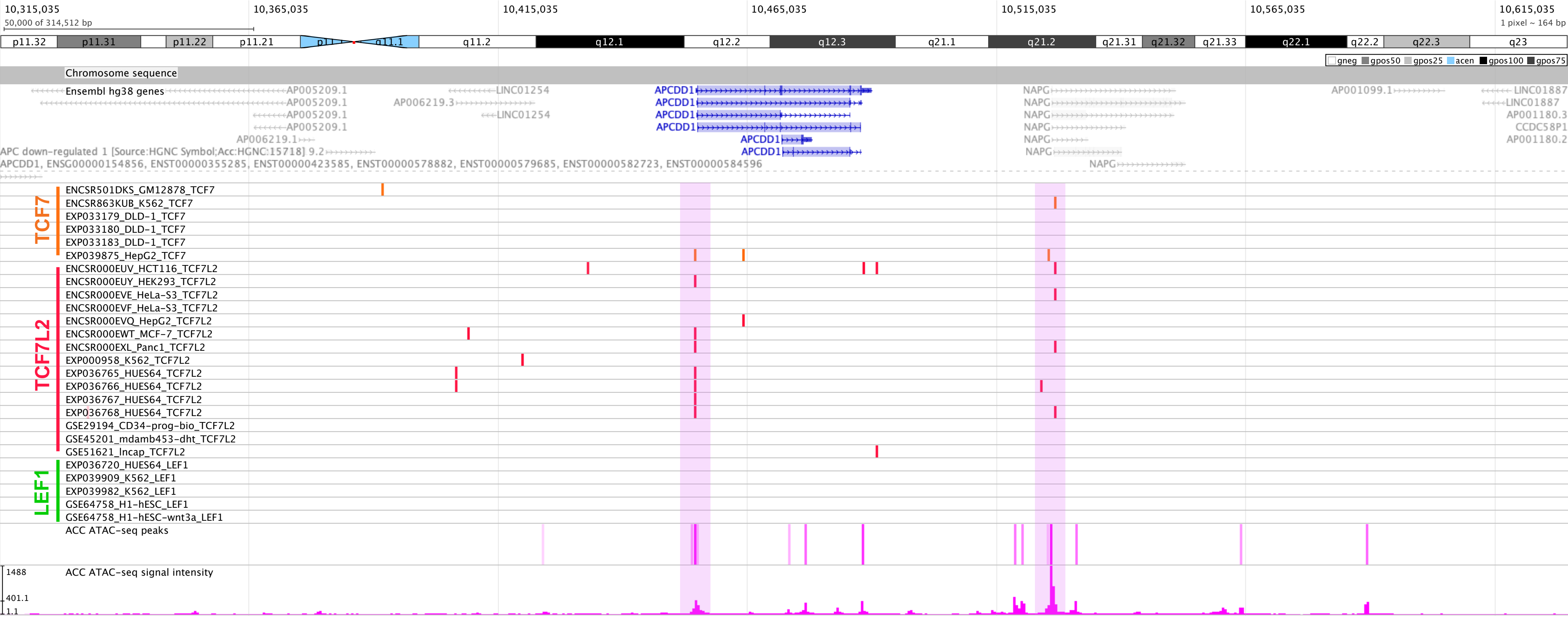

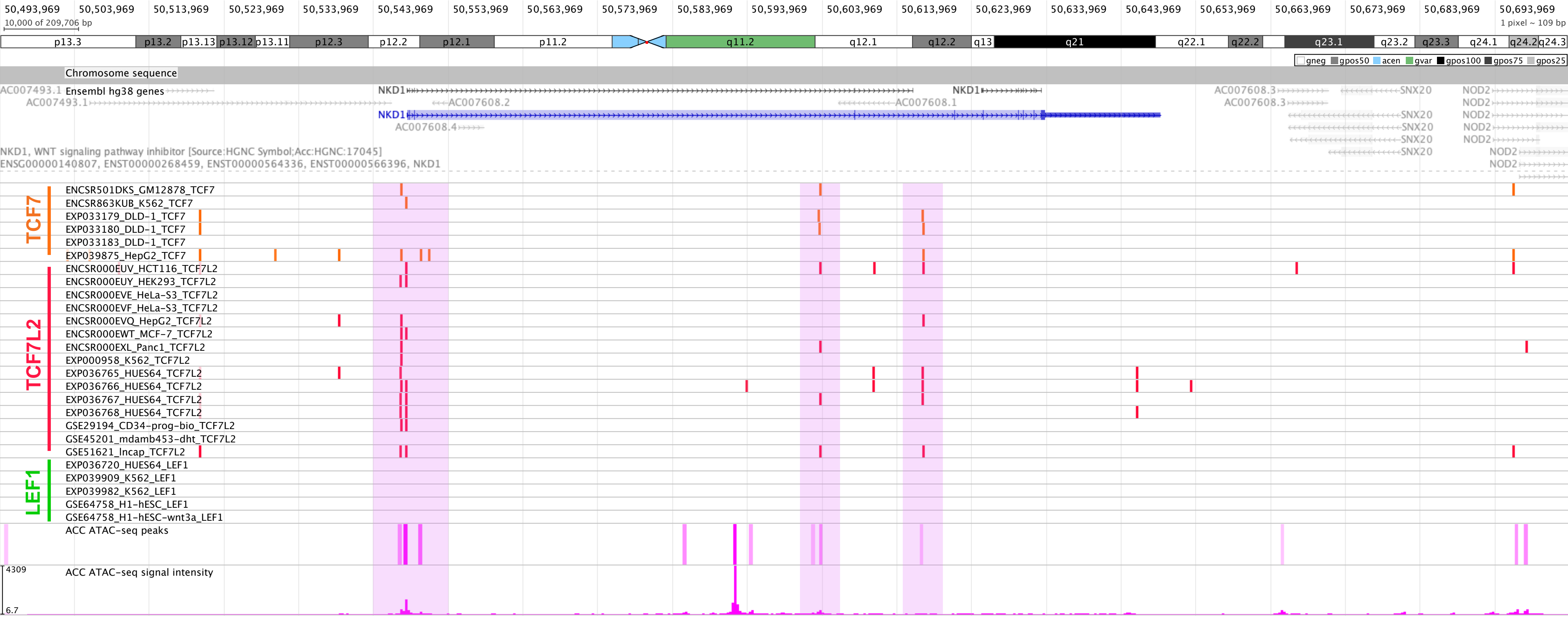

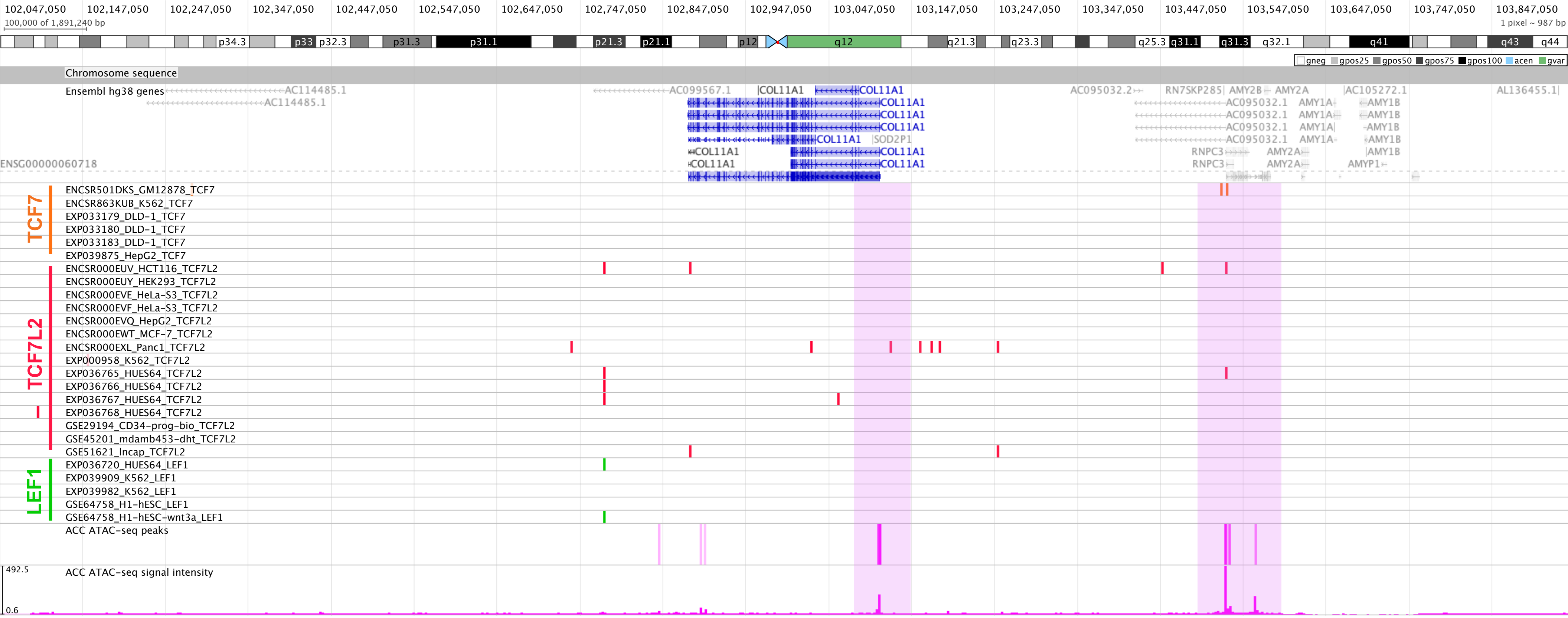

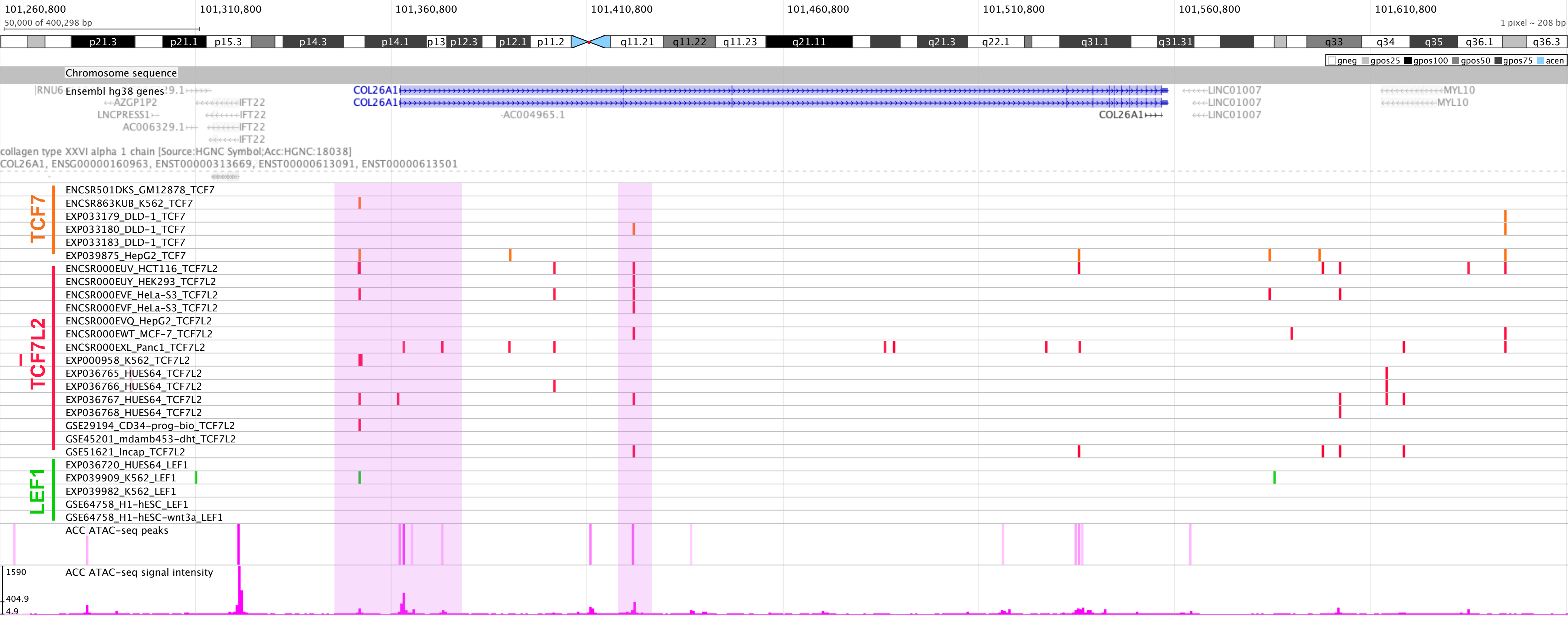

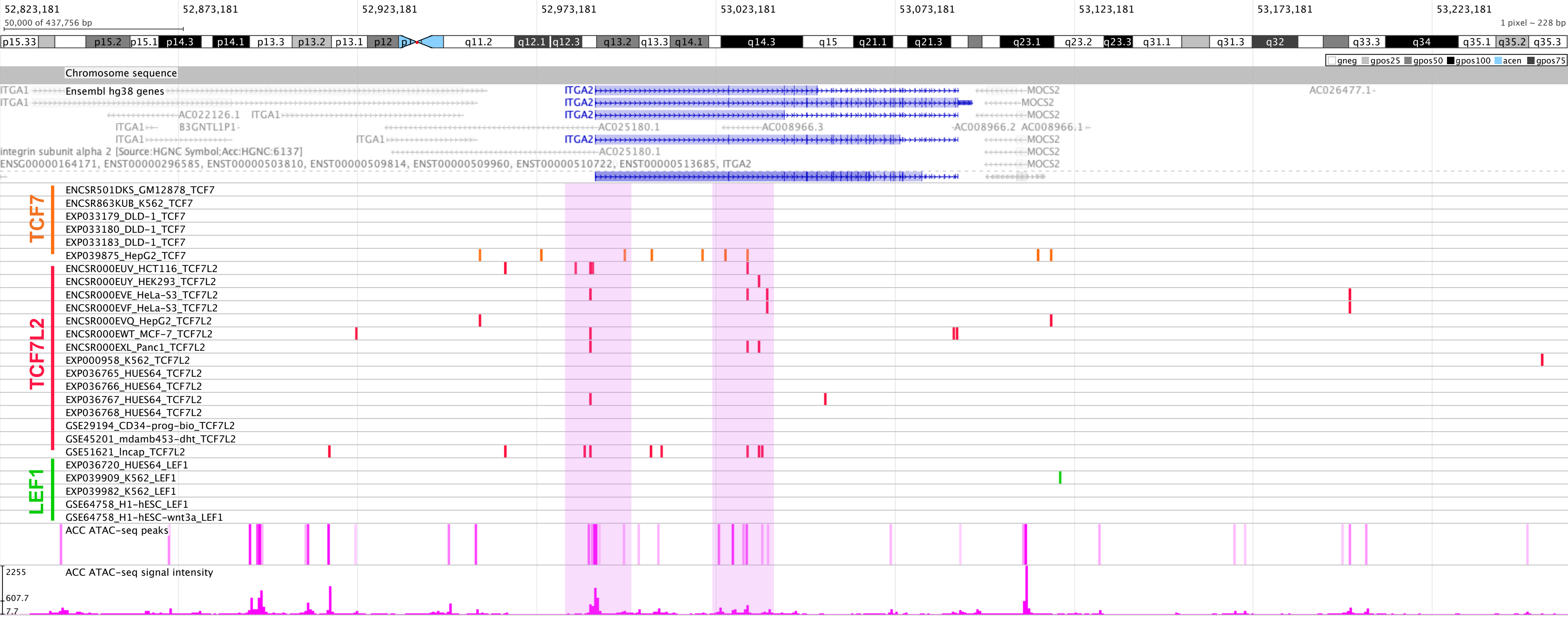

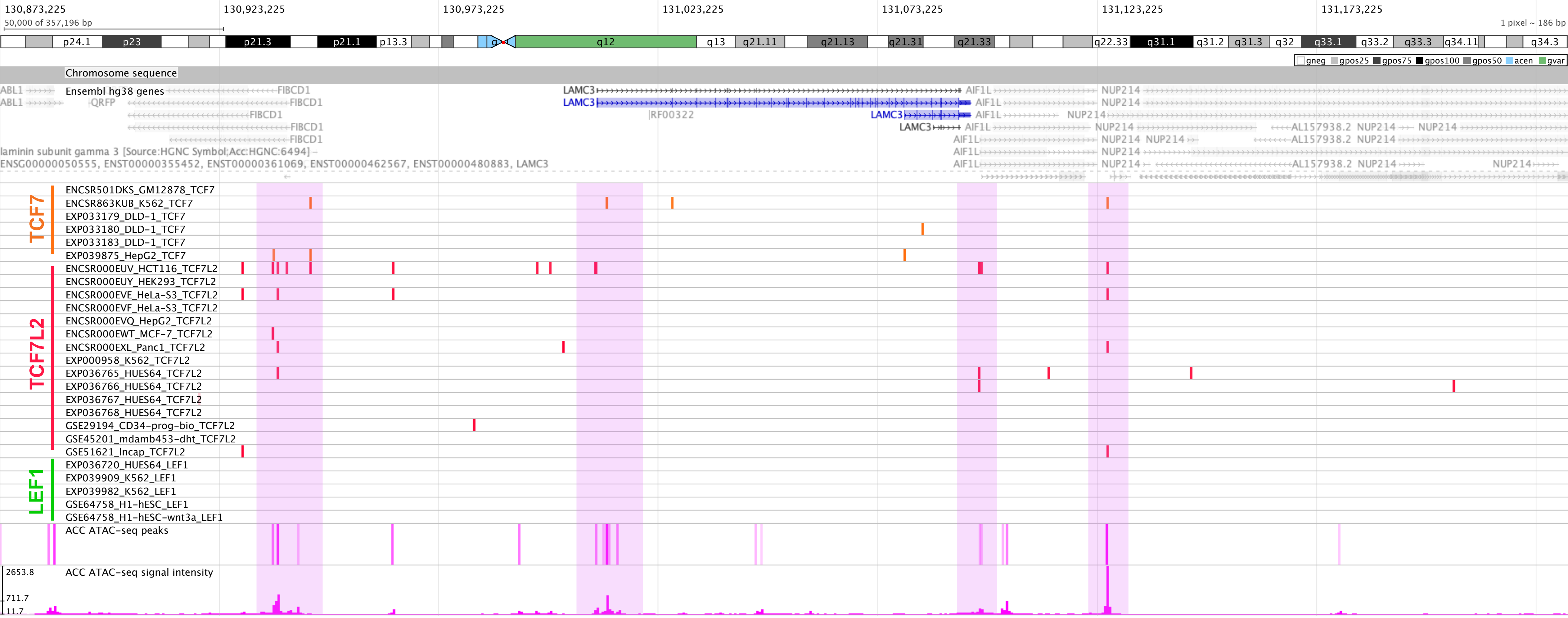

### Supplementary Figure 2

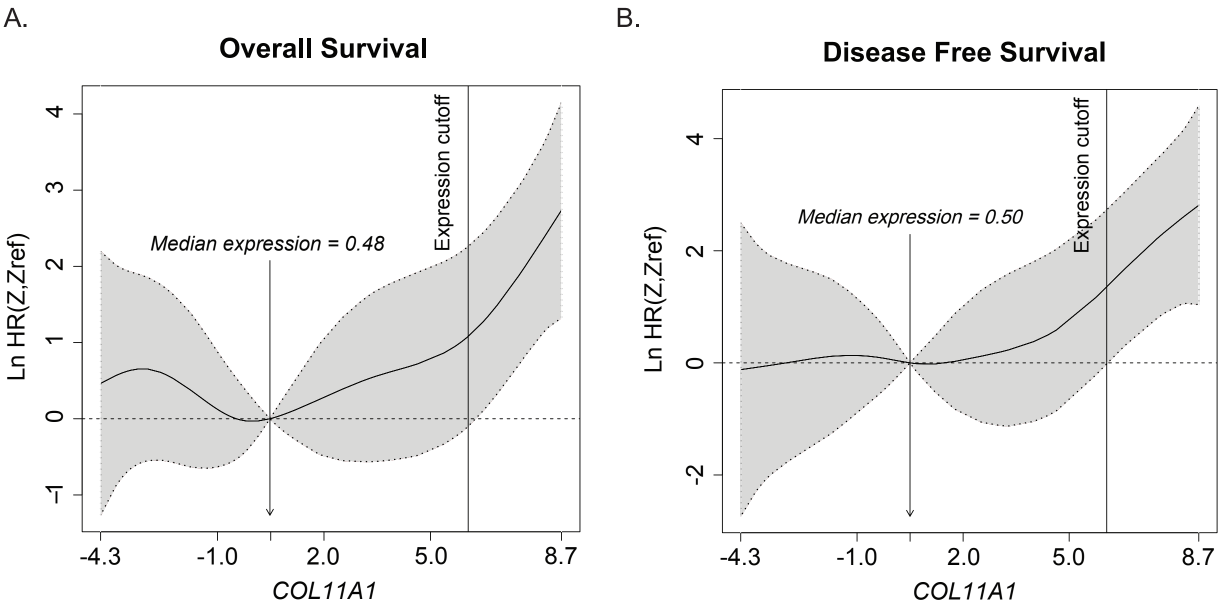

### Supplementary Figure 3

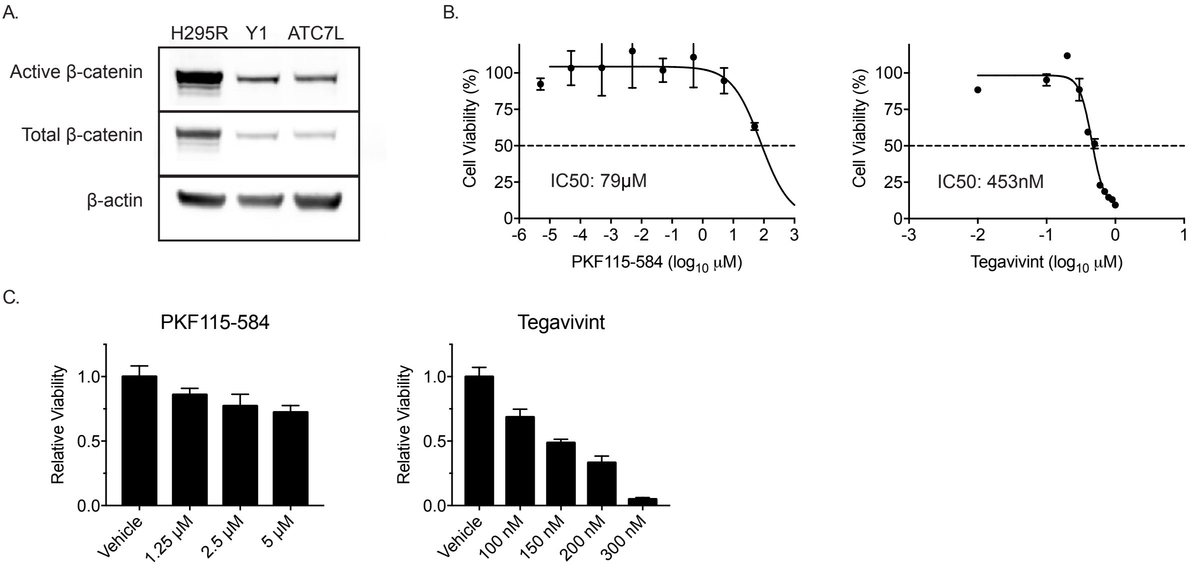

### Supplementary Figure 4

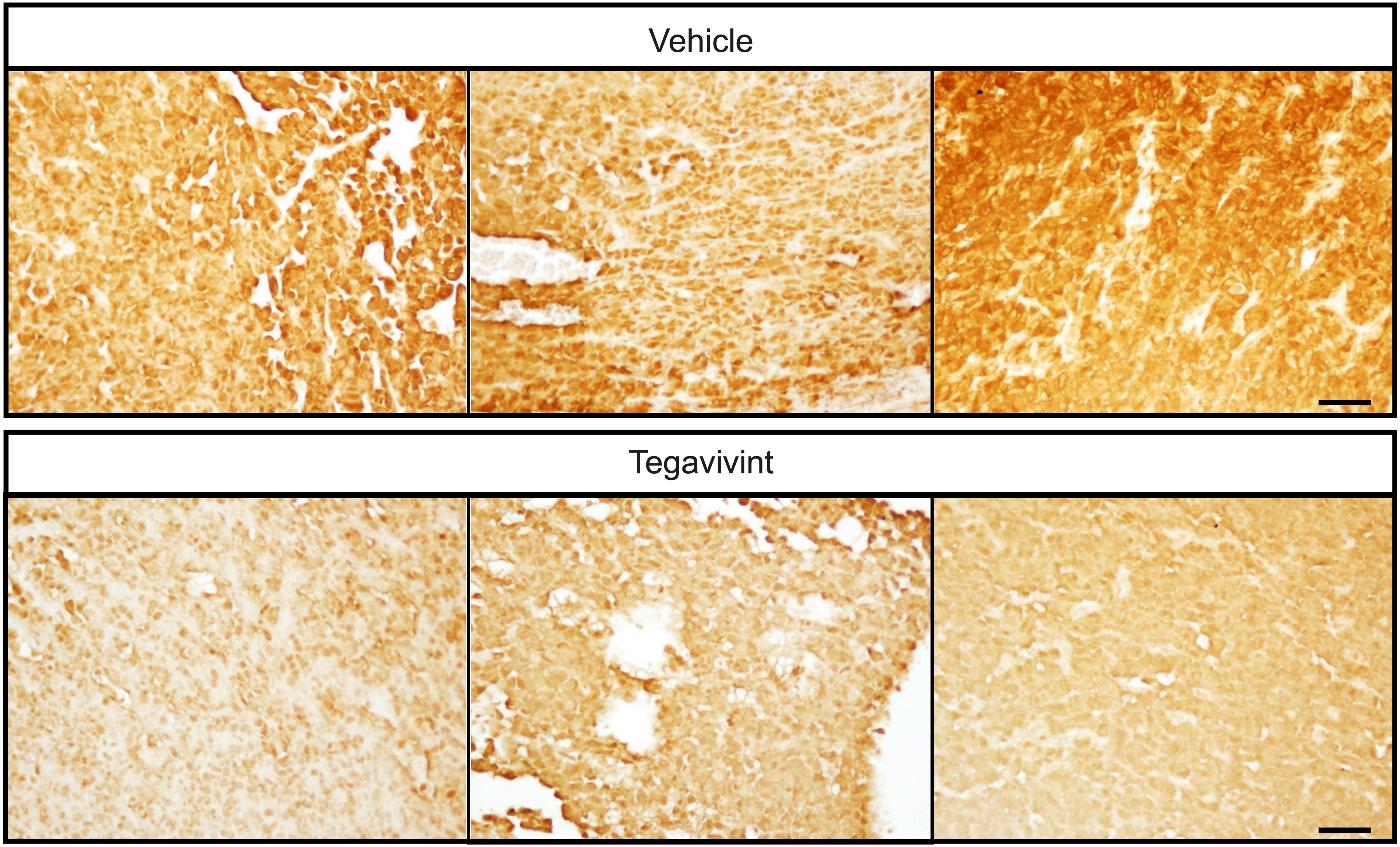
