## Supplementary Table 2 for "Targeting oncogenic Wnt/β-catenin signaling in adrenocortical carcinoma disrupts ECM expression and impairs tumor growth"

| Gene | Forward Primer 5' -> 3' | Reverse Primer 5' -> 3' |
|---|---|---|
| *HPRT1* | TGACACTGGCAAAACAATGCA | GGTCCTTTTCACCAGCAAGCT |
| *AXIN2* | AAGTGCAAACTTTCGCCAAC | ACAGGATCGCTCCTCTTGAA |
| *LEF1* | CTTTATCCAGGCTGGTCTGC | TCGTTTTCCACCATGTTTCA |
| *APCDD1* | ATGCCACCCAGAGGATGTTC | GATGGTCAGGTCTGCCTTTG |
| *COL11A1* | GACTATCCCCTCTTCAGAACTG | CTTCTATCAAGTGGTTTCGTGGTTT |
| *COL26A1* | CAGCAGCTGAGAGAGGCCCT | GCCACCCCTCTTCATCTTGAG |
| *ITAG2* | AACTCTTTGGATTTGCGTGTG | TGGCAGTCTCAGAATAGGCTTC |
| *LAMC3* | GCTCCGAGGAATGCACGTT | TGTCATCGCACTGGAGGTGTA |
